# A CANCER PERSISTENT DNA REPAIR CIRCUIT DRIVEN BY MDM2, MDM4 (MDMX), AND MUTANT P53 FOR RECRUITMENT OF MDC1 AND 53BP1 TO CHROMATIN

**DOI:** 10.1101/2024.01.20.576487

**Authors:** Viola Ellison, Alla Polotskaia, Gu Xiao, Pamella Leybengrub, Weigang Qiu, Rusia Lee, Ronald Hendrickson, Wenwei Hu, Jill Bargonetti

## Abstract

The influence of the metastasis promoting proteins mutant p53 (mtp53) and MDM2 on ***C**ancer **P**ersistent **R**epair* (CPR) to promote cancer cell survival is understudied. Interactions between the DNA repair choice protein 53BP1 and wild type tumor suppressor protein p53 (wtp53) regulates cell cycle control. Cancer cells often express elevated levels of transcriptionally inactive missense mutant p53 (mtp53) that interacts with MDM2 and MDM4/MDMX (herein called MDMX). The ability of mtp53 to maintain a 53BP1 interaction while in the context of interactions with MDM2 and MDMX has not been described. We asked if MDM2 regulates chromatin-based phosphorylation events in the context of mtp53 by comparing the chromatin of T47D breast cancer cells with and without MDM2 in a phospho-peptide stable isotope labeling in cell culture (SILAC) screen. We found reduced phospho-53BP1 chromatin association, which we confirmed by chromatin fractionation and immunofluorescence in multiple breast cancer cell lines. We used the Proximity Ligation Assay (PLA) in breast cancer cell lines and detected 53BP1 in close proximity to mtp53, MDM2, and the DNA repair protein MDC1. Through disruption of the mtp53-MDM2 interaction, by either Nutlin 3a or a mtp53 R273H C-terminal deletion, we uncovered that mtp53 was required for MDM2-53BP1 interaction foci. Our data suggests that mtp53 works with MDM2 and 53BP1 to promote CPR and cell survival.

## Introduction

In breast cancer, often the *TP53* gene contains missense mutations and the *MDM2* gene is over expressed [1]. The proteins produced by these altered alleles promote aggressive cancers and metastasis [2, 3]. However, the normal *TP53* gene encodes the site-specific DNA binding transcription factor p53, which is a multifunctional genome guardian that is the most highly mutated gene in cancers [4, 5]. The wild-type (wtp53) protein has evolutionarily conserved transcription activation functions and genome maintenance to promote DNA damage recognition, signaling, and repair [4, 6]. While wtp53 is conserved *from C. elegans* to humans for the transcriptional activation of cell cycle inhibitors and apoptosis inducers, only in vertebrate cells does wtp53 activate the *MDM2* gene to establish the equilibrium of p53 levels and promote attenuation of the stress response; there is no identified *MDM2* homolog in either *Drosophila or C. elegans* [7, 8]. In response to a variety of cellular stresses, particularly those which compromise genome integrity, wtp53 and MDM2 crosstalk to allow for wtp53 to induce expression of hundreds of genes that collectively determine one of three broad cell fate outcomes, which are cell cycle arrest, senescence, or cell death [9]. The MDM2/MDMX (also called MDM4) E3 ubiquitin ligase complex targets wtp53 for ubiquitin-mediated proteolysis [9]. MDM2 and MDMX are RING domain E3 family members that contain N-terminal and C-terminal regions that bind, and regulate, wtp53 and many other proteins [10]. MDM2 forms a homodimer that is competent to ubiquitinate wtp53, but the MDMX protein lacks intrinsic ubiquitin ligase activity; MDMX when in a heterodimer with MDM2 is important for E2 recruitment and enhances MDM2 specific E3 ubiquitin ligase activity [11]. Both MDM2 and MDMX have been shown to interact with proteins that are part of DNA repair pathways [12]. While wtp53 is well known for its interaction with the DNA repair protein 53BP1, only a few studies have examined the interaction of mtp53 with 53BP1 and none in the context of changes to MDM2 and MDMX in cancer cells [13–17].

Cancer cells often overexpress stable mtp53 protein that has a loss of function (LOF) caused by missense mutations that disrupt site-specific DNA binding activity and a subset of these mtp53 proteins possess gain-of function (GOF) [18, 19]. Importantly, these mtp53 proteins retain the ability to non-specifically bind DNA, maintain protein-protein interactions with many partners, and also gain new protein-protein interaction partners [19–21]. A quintessential property of GOF mtp53 in cancer cells is that it is maintained at stable high cellular levels despite the ability of mtp53 to form complexes with MDM2 [22–24]. Moreover, some splice variants of MDM2 promote mtp53 GOF [25]. These observations suggest mtp53 works with MDM2 and MDMX to drive functions that promote tumorigenesis [26]. Examples of mtp53 GOF activities that suggest alterations in DNA metabolism promoting cell survival have been reported by many laboratories, and include mtp53-PARP1 modulation of DNA replication and repair intermediates [27–29], mtp53-Treslin modulation of replication fork assembly [30], and the observation that inhibition of PARP or Treslin or MDM-Two Binding Protein (MTBP) leads to DNA damage [31]. Both MDM2 and MDMX also participate in regulating DNA replication [32, 33].

Here we describe the usage of a SILAC phospho-proteome screen performed in T47D breast cancer cells that express mtp53 (L194F) to identify changes in the chromatin residence of proteins in cells that lack MDM2, which we posited would reveal MDM2-MDMX-driven tumorigenesis mediating components. We found reduced recovery of several proteins involved in shaping the nuclear landscape, including the chromosome protection and DNA repair proteins p53 Binding Protein 1 (53BP1) and Mediator of DNA damage Checkpoint 1 (MDC1). Using a combination of *in situ* techniques (immunofluorescence and the Proximity Ligation Assay (PLA)) and *in vitro* techniques (including chromatin fractionation and immunoprecipitation) we observed a hierarchy of protein-protein interactions that were dependent on both mtp53 and MDM2. We observed a dependence on mtp53 for the MDM2-53BP1 interaction. The interaction between 53BP1 and MDC1 is required for 53BP1 recruitment to stressed DNA in the absence of exogenous DNA damage through a Proline-Serine-Threonine repeat (PST) domain [34, 35], and we observed significantly reduced 53BP1-MDC1 foci in the absence of MDM2 protein, or when the MDM2-mtp53 interaction was inhibited using the small molecule inhibitor Nutlin 3a. These observations suggest that in the context of mtp53, the mtp53-MDM2-MDMX complex actively participates in regulating recruitment of 53BP1 to sites of DNA replication stress and endogenous DNA damage. We propose the function of mtp53 working together with 53BP1 and MDM2 potentiates what we are calling ***C****ancer **P**ersistent **R**epair* (CPR) to keep tumor cells viable. The 53BP1 promotes NHEJ error prone repair and thus would contribute to the observed GOF mtp53 promoted genomic instability and metastasis [36].

## Materials and Methods

### Chemicals and Antibodies

Solvents and standard chemicals for buffers were obtained from Sigma-Aldrich (St. Louis, MO, USA) and Fisher Scientific unless otherwise indicated. Antibodies used for western blotting (*WB*), immunofluorescence staining (*IF*), immunoprecipitation (*IP*), and Proximity Ligation Assay (*PLA*) were purchased from the following (usage denoted in italics): (1) rabbit p53 Sigma cat# A300-247A (*PLA*), and Proteintech cat# 10442-1-AP (*WB*); (2) mouse p53 DO1 Santa Cruz Biotechnology cat# sc-126 (*PLA, WB*); (3) mouse p53 DO1-HRP Santa Cruz Biotechnology cat# sc-126 HRP (*WB*); (4) rabbit MDMX Proteintech cat# 17914-1-AP (*WB*); (5) rabbit MDM2 R&D Systems cat# AF1244 (*WB*); (6) rabbit 53BP1 Cell Signaling Technology cat# 4937 (*WB, IF*); (7) rabbit phospho-Serine 1778 53BP1 Cell Signaling Technology cat# 2675 (*WB, IF*); (8) rabbit phospho-Serine 25 53BP1 Sigma cat# PLA 0126 (*WB, IF PLA*); (9) rabbit MDC1 Sigma cat# PLA0016 (*WB, IF, PLA*); (10) rabbit MCM4 Cell Signaling Technology cat# 12973 (*WB*); (11) mouse Actin-HRP Sigma cat# A3854 (*WB*); (12) mouse Lamin A cat# SAB4200420 (*WB*); (13) mouse PARP1 BD Biosciences cat# 51-6639GR (*WB*) ; (14) goat 53BP1 Sigma cat# PLA0303 (*PLA*); (15) goat anti-mouse HRP Sigma cat# A3682 (*WB*); (16) goat anti-rabbit Proteintech cat# SA00001-2 (*WB*); (17) mouse Cyclin A Santa Cruz Biotechnology cat# sc-271682 (*WB*); (18) rabbit Cyclin A Cell Signaling Technology cat# 67955S (*IF*); (19) mouse Cyclin B Santa Cruz Biotechnology cat# sc-245 (*WB*); (20) rabbit p21 Cell Signaling Technology cat# 2947S (*WB*); (21) γH2AX phospho-Ser139 Cell Signaling Technology cat# 9718S (*WB, IF*); (22) rabbit Poly ADP Ribose Cell Signaling Technology cat# 83732S (*WB*); (23) mouse MDM2 SMP14 Santa Cruz Biotechnology cat# sc-965 (*IP*); (24) mouse IgG Santa Cruz Biotechnology cat# sc-2025 (*IP*); (25) Purified mouse MDM2 4B2 [37] and (26) purified mouse MDM2 2A9 [37] were used for PLA and IP and prepared as described [38]. Nutlin3a (cat# S8059), Talazoparib (cat# S7048), Temozolomide (cat# S-1237) were purchased from Selleckchem, Etoposide from Sigma (cat# E1383) and Aphidicolin from ApexBio (B7832). Stock solutions for drugs were prepared in 100% sterile DMSO (G-Biosciences cat# 786-1388).

### Cell culture and drug treatments

All cell lines-MDA-MB-468 (ATCC), MDA-MB-468 CRISPR clone G6 (R273H*fs*347Δ360-393) [39], MDA-MB-231 (ATCC), MCF7 (ATCC), T47D mlp [40], T47D mlp*shmdm2* [40], T47D mlp*shmdmx* [40], HCT116 *p53+/+* and HCT116 *p53-/-* (gift from Bert Vogelstein) were cultured in complete media (Dulbecco’s Modified Eagle Medium with 4.5g/L glucose or McCoy’s (HCT116 *p53+/+* and HCT116 *p53-/-*) containing 10% fetal bovine serum, 50μg/ml penicillin/streptomycin) at 37°C with 5% CO_2_ and passaged by trypsinization and dilution. For all drug treatments, culture medium on cell populations was removed and fresh medium containing either the working concentration of agent or vehicle (DMSO or H_2_O) was added, followed by incubation of cells for the indicated time. For enrichment of early S-phase and G2/M populations, cells were grown in 5µM Aphidicolin for 24hr, and then either harvested (Aph 24.0), or incubated for an additional 10hr (Aph 24.10) after washing with PBS and addition of fresh media.

### Flow cytometry

Cultures were harvested by trypsinization, collected by centrifugation (500 x *g*), washed with ice cold PBS twice, and then resuspended in ice-cold PBS at a density of 1x10^6^ cells/ml. Cells were fixed in a final concentration of 70% Ethanol by adding the cell suspension dropwise into a solution of 100% ethanol while vortexing continuously. Fixed cells were stored at -20°C for 24 hours, and then collected by centrifugation, washed twice with ice-cold PBS and then resuspended and incubated at 37°C for 15 min in 500μl Propidium Iodide staining solution (0.1% Triton X-100, 200μg/ml RNAse A, 60 μM Propidium Iodide in PBS). Following staining, cell samples were filtered through a nylon mesh into polystyrene tubes, and then analyzed on a BD^TM^ FACSCalibur instrument. Minimally 10,000 events were counted for each sample and acquired data was analyzed using BD^TM^ CellQuest or FlowJo software.

### Immunofluorescence staining (IF)

Cells were cultured on 12-well uncoated plates (MatTek, Cat# P12G-1.5-14-F) until ∼60% confluency, washed 3x with cold PBS (1 min/wash) and then fixed at room temperature (r.t.) with 4% Paraformaldehyde in PBS for 15 min. Post-fixation, cells were washed 3x with 1ml cold PBS (5 min/wash), permeabilized at r.t. for 15 min with cold IF Permeabilization Buffer (0.5% Triton X-100 in PBS), blocked at r.t. for 1 hr with IF Blocking Buffer (5% Normal Goat Serum, 0.2% Triton X-100 in PBS), and then incubated at r.t. for 2 hr with primary antibodies (1:1000 53BP1; 1:3000 53BP1^ser25^; 1:1000 53BP1^ser1778^; 1:1000 Cyclin A2). After incubation with primary antibodies, cells were washed 3x with cold PBS, and then incubated at r.t. for 1hr with the secondary antibody in the dark (anti-rabbit Alexa Fluor 594; ThermoFisher Cat #11012). After washing with PBS as described above, cells were stained for 5 min with 10μM Hoescht 33342 in PBS, washed at r.t. 1 min with PBS, and then allowed to dry overnight at 4°C before mounting using CitiFluor AF1 (50µl/well; cat# 17970-25). Images were taken using the Nikon A1 confocal microscope and processed with the Nikon NIS Element software, ImageJ and Cell Profiler. Data analysis including appropriate statistical tests (ANOVA, Kruskal-Wallis or Mann-Whitney U) were performed using GraphPad Prism 9.

### EdU labeling for measurement of S-phase population

For measurement of the S-phase population, cells grown in 12-well glass bottom plates were incubated with 20µM EdU added directly to the culture medium (Invitrogen; cat# A10044) for 15 min. Following fixation and permeabilization as specified in the immunofluorescence protocol, cells were washed twice with ice cold 3%BSA in PBS for 5 min, once with PBS for 5 min, and then incubated in the dark at r.t. for 30 min with either Alexa Fluor 594 or 647-Azide (Invitrogen; A10270 and A10277 respectively) under click chemistry reaction conditions (100µl/well 4µM Azide compound, 1mM Copper Sulfate (Sigma; cat# 451657), 5mM Sodium Ascorbate (freshly prepared, Sigma; cat# A7631), 50mM Tris, 150mM NaCl, final pH 7.5). After the incubation, the reaction buffer was discarded and cells were washed (1ml/well, 5 min/wash) once with PBS, once with 3% BSA in PBS, once with PBS and then blocked and subjected to immunofluorescence staining for specific antigens; all steps were performed in the dark.

### Proximity Ligation Assay (PLA)

The protocol for the proximity ligation assay was performed as described previously [29] using the Sigma Aldrich Duolink System^TM^ (anti-mouse, rabbit or goat (-) or (+) secondary antibodies; either 594 (cat# DUO92008) or 647 (cat# DUO92013) detection kit) with modifications indicated below (unless indicated otherwise, all incubations were performed as specific by the manufacturer). Cells were seeded in 12-well un-coated glass-bottomed plates (MatTek) at 1x10^5^ cells per well in complete media, cultured until 50% confluency and then washed 3x with ice cold PBS for 1 min/wash followed by fixation with 4% Paraformaldehyde in PBS for 15 min. Next, cells were permeabilized with ice cold PBS containing 0.5%Triton X-100 for 15 min, blocked, and then incubated with primary antibodies (55 μl/well) for 2 hr using the indicated dilutions (prepared in Duolink Ab diluent): (1) goat anti 53BP1 1:1000; (2) rabbit anti p53 Sigma 1:2000; (3) rabbit anti MDC1 1:2000; (4) mouse anti MDM2 4B2 1:500 (final concentration 1μg/ml) or 2A9 1:1000 (final concentration 1μg/ml). After washing 3x (1ml/wash), cells were incubated with the appropriate PLA Plus/Minus pair of secondary antibody probes (55 μl/well), and then subjected to the ligation and amplification steps. After the amplification step cells were washed 2x with Buffer B as indicated, 1x with 0.01x Buffer B containing 1μg/ml Hoescht 333243 for 5 min, once with 0.01x Buffer B for 1 min and then air dried in the dark at 4°C after first removing remaining wash buffer by pipetting. Cells were mounted as described in the immunofluorescence protocol and images were taken using the Nikon A1 confocal microscope and processed with the Nikon NIS Element software, ImageJ and Cell Profiler. Data analysis including statistical tests (Kruskal-Wallis or Mann-Whitney U) were performed using GraphPad Prism 9.[40]

### Cell extract preparation, SDS-PAGE, and western blot analysis

Total cell lysates were prepared using Phosphate/Chaps (PC) Buffer (50mM KPO_4_, 1mM DTT, 1mM EDTA, 50mM NaF, 50μM NaV, and 10mM β-glycerolphosphate, 7mM CHAPS, 10% glycerol, 350mM NaCl, pH 7.4) supplemented with Pierce Complete^TM^ protease inhibitor cocktail tablet (ThermoFisher Cat# A32961) as specified by the manufacturer. Sub-confluent cell cultures from 6 cm plates were harvested by either trypsinization or scraping using a rubber policeman, collected by low-speed centrifugation, washed by resuspending in 5ml cold PBS, re-pelleted and then resuspended in ice-cold PC Buffer (1x10^6^ cells/100μl buffer). Resuspended cells were incubated on ice for 30 min and cell lysates were clarified by centrifugation at 10,000 rpm at 4°C for 30 min in an Eppendorf 5415R microfuge. After determining the protein concentration of each sample using the Bio-Rad Bradford assay, aliquots (10-25μg) of cell lysates were prepared for SDS-PAGE by mixing with an equal volume of 2x NuPAGE^TM^ sample buffer containing 50mM DTT, heated at 75°C for 10 min, and then incubated at r.t. for 10 min after addition of Iodoacetamide to a final concentration of 50mM to inhibit disulfide bond formation. Samples were then loaded onto an appropriate concentration SDS-PAGE gel (typically 8% gels; 30% Acrylamide:0.4% Bis-acrylamide) for western blot analysis (electroblotting performed onto nitrocellulose membrane). Blots were blocked using 5% Non-Fat Dry Milk Buffer (NFDM Buffer; 5% NFDM in 10mM Tris pH 7.4, 150mM NaCl), and all primary antibodies were diluted 1:1000 in 1% NFDM Buffer containing 0.1% Triton X-100 except for the following: (1) rabbit p53 Proteintech was used at 1:10,000-1:20,000; (2) rabbit MDM2 R&D Systems was diluted in 3% BSA in PBS containing 0.1% Triton X-100. Primary antibodies were detected by chemiluminescence using the appropriate HRP-conjugated anti-rabbit or anti-mouse secondary antibody and the Pierce Super Signal^™^ West Pico detection system.

### MDM2 and mtp53 co-immunoprecipitation from MDA-MB-468 and MDA-MB-468 CRISPR clone G6 (R273Hfs347Δ360-393)

Mouse IgG (Santa Cruz Biotechnology sc-2025)-protein A/G agarose, 4B2-protein A/G agarose and SMP14-protein A/G agarose were prepared in siliconized 1.5ml microfuge tubes by incubating 60μg antibody with 240μl Protein A/G agarose beads (Fisher Scientific cat# IP1010ML) in a final volume of 1.2 ml of 1M NaCl, 0.1M Na Borate pH 8.5 for ≥3hr at 4°C. After binding, antibody/beads were allowed to settle by gravity then centrifuged 1 min 1500 rpm at 4°C. After removal of the supernatant, the antibody beads were successively washed (10 min/wash) with (1) 2x with 1M NaCl, 0.1M Na Borate pH 8.5, (2) 2x with PBS, (3) 1x with 1mg/ml BSA in PBS, and (4) 2x PBS. Following the last wash, the antibody beads were resuspended in 720μl PBS (25% slurry); 20μl of antibody beads (5μg antibody) was used for each IP reaction. Using the procedure for extract preparation above, one sub-confluent 15cm plate (∼1x10^7^ cells) of each cell line was harvested, washed 2x (20ml/wash) with ice-cold PBS, and the cell pellets were resuspended in 1.6ml of IP Buffer (25mM Tris pH 7.5, 1mM DTT, 1mM EDTA, 50mM NaF, 50μM NaV, and 10mM β-glycerolphosphate, 7mM CHAPS, 10% glycerol, 200mM NaCl, complete protease inhibitor cocktail). Cell lysates were incubated on ice for 30 min, clarified by centrifugation at 10,000 rpm at 4°C for 30 min in an Eppendorf 5415R microfuge, and assayed for protein content using the Bio-Rad Bradford Assay. For each extract 1-1.5mg of total protein was used in each IP reaction (set up in siliconized 1.5ml microfuge tubes), which were incubated typically overnight (≤12hr) at 4°C (cold room) with rotation (Labquake; Barnstead Thermolyne). After the incubation, antibody-beads were allowed to settle by gravity, the supernatant removed, 1.4ml of IP Buffer was added and IP samples washed with rotation in the cold room for 15 min. This wash step was repeated two more times with IP sample collection by centrifugation (4°C 2 min, 2000 rpm) instead of gravity, and after removal of the last wash, IP samples were resuspended in 54μl 1x NuPAGE containing 25mM DTT, heated at 75°C for 10 min and then incubated at r.t. for 10 min after addition of Iodoacetamide (6μl). IP samples (12.5% or 25% of each) were subjected to western blot analysis.

### MTT Assay

T47D cells seeded in 12 well plates at confluency of 10% were grown to 50% confluency, treated (in triplicate) with the indicated drug for 24hr (10μM Nutlin3a or 50μM Etoposide), washed with 2ml/well of PBS after drug removal, and then incubated with 1ml fresh complete media containing 0.5mg/ml MTT (3-(44,5-dimethylthiazol-2-yl)-2,5-diphenyltetrazoliumbromide) at 37°C for 30 min. After the incubation, cells were lysed and the Formazan product of MTT metabolism solubilized by addition of 1ml stop solution (1% Triton X-100, 104mM HCl in isopropanol). Product formation was measured using a microtiter plate reader (absorbance at λ_550_-λ_620_). Background (defined by wells with no cells) was subtracted and the data from each cell line was normalized to the negative control (untreated cell population).

### Chromatin Fractionation Assay

Chromatin localization of proteins was measured using the chromatin fractionation assay. Briefly, cell populations (one 50-60% confluent 10 cm plate/sample) were harvested using a rubber policeman, pelleted and washed with ice-cold PBS, and then resuspended in Buffer A (10mM HEPES, 10mM KCl, 1.5mM MgCl_2_, 300mM Sucrose, 1mM DTT, 10% Glycerol, 0.1mM PMSF, 1μg/ml Leupeptin, 1μg/ml Pepstatin A, and 2μg/ml Aprotinin) in a volume of 300μl (cell density ∼1x 10^7^ cells/ml). After hypotonic swelling, cells were then lysed by addition of Triton X-100 (final concentration 0.15%) on ice for 15 min, and the nuclei pelleted by low-speed centrifugation (4 min at 1,300 x *g* at 4℃; supernatant (S1) saved as cytosolic extract). After washing twice with 300 µl Buffer A + 0.15% Triton X-100, nuclei were lysed by resuspending in 300 µl of Buffer B (3mM EDTA, 0.2mM EGTA, 1mM DTT, 0.1mM PMSF, 1μg/ml Leupeptin, 1μg/ml Pepstatin A, and 2μg/ml Aprotinin) and incubating on ice for 30 min. Chromatin from each sample was pelleted by centrifugation (1,700 *x g* at 4℃ for 4 min), washed with 500 µl of Buffer B twice, and then resuspended in 300 µl of Buffer B and sheared by sonication. The protein concentration of both S1 and chromatin fractions were quantified using the Bio-Rad Bradford assay and subjected to SDS-PAGE and western blot analysis (7.5 µg of the S1 fraction;10 µg of the chromatin fraction).

## Results

### Loss of MDM2 influences the chromatin binding activity of 53BP1

To identify the MDM2 dependent and mtp53 components of the breast cancer cell proliferation and metastasis signaling pathways we used our previously developed shRNA cell line tools for the comparison of outcomes with, and without, the knockdown of MDM2 and MDMX [40–42]. We previously showed that loss of MDM2 impairs T47D and MCF-7 cell proliferation [40, 42, 43]. Additionally, we demonstrated that loss of MDM2 inhibits triple negative breast cancer MDA-MB-231 and MDA-MB-468 cell metastasis [40, 41]. We performed stable isotope labeling with amino acids in cell culture (SILAC) in T47D control and T47D sh*mdm2* (MDM2-depleted) cells. We purified chromatin-associated phospho-peptides, which were then subjected to LC-MS/MS mass spectrometry analysis (see Fig. 1A for workflow). The analysis revealed 1381 peptides corresponding to 317 unique proteins under-represented on chromatin derived from MDM2-depleted cells and 20 proteins over-represented (Fig. 1A and http://borreliabase.org/~wgqiu/clickme-khalikuz/temp-Points.html). Prominent among the 1381 peptides identified were those for the DNA repair proteins 53BP1 and MDC1. Both 53BP1 and MDC1 are central components of chromatin localized factors that are mobilized early in the response to double-strand breaks (DSBs) to delineate damaged chromatin domains marked by phosphorylated and ubiquitinated forms of H2AX [44–46]. Given that 53BP1 participates in a signaling axis with p53 [15], and MDM2 interacts with MDC1 (also called NFBD1) [47], we then asked if the MDM2 expression level influenced 53BP1 recruitment to chromatin.

**Figure 1:**
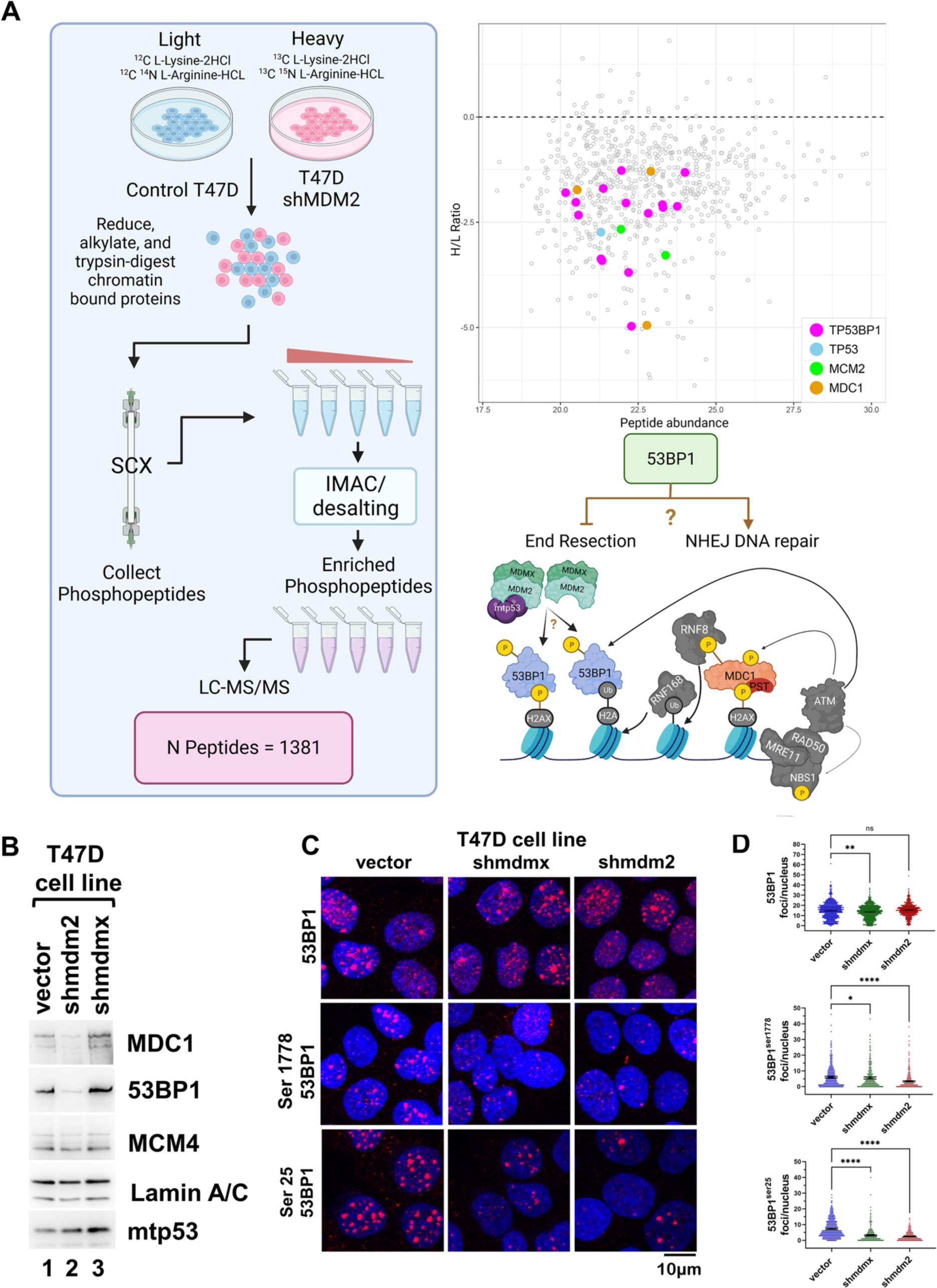
Reduced MDM2 protein in T47D cells causes reduced chromatin phosphoproteins 53BP1 and MDC1. **(A)** The workflow of the SILAC analysis is shown with a diagram made on BioRender. Chromatin isolated from a mixture of T47D vector control cells (MDM2-competent) cultured in natural amino acid medium, and T47D*shmdm2* cells (MDM2-depleted) cultured in heavy isotope amino acid medium, was subjected to proteolysis followed by phospho-peptide purification and enrichment. Scatter plot represents the H/L ratio versus abundance of peptides identified by mass-spectrometry, with those corresponding to TP53BP1 (magenta), TP53 (blue), MCM2 (green), and MDC1(brown) highlighted. Cartoon depicts known chromatin-based events that contribute to recruitment of 53BP1 to DNA double strand breaks (DSBs). 53BP1 DNA end binding promotes NHEJ factor assembly. **(B)** Chromatin (5µg) isolated from T47D vector control, T47D*shmdm2*, and T47D*shmdmx* cells was subjected to SDS-PAGE/western blot analysis for 53BP1, MDC1, MCM4, Lamin A/C, and mtp53. **(C and D)** IF of total 53BP1, phospho-53BP1^ser25^ or phospho-53BP1^ser1778^ in T47D vector control, T47D*shmdm2*, and T47D*shmdmx* cell lines. Primary antibodies were detected with anti-rabbit Alexa Fluor 594; nuclei were stained with Hoescht 33342. Confocal images for six fields for each were acquired and the number of 53BP1 foci/nucleus from each cell population (>300 nuclei) was determined using Cell Profiler. Data representations are mean foci/nucleus with 95% confidence interval from one representative experiment (n=3); graphs were prepared and statistical analysis (Kruskal-Wallis) performed using Prism; p < .05 (*), p < .01 (**), p < .001 (***), p < .0001 (****), p > .05 (ns).

We validated that decreased MDM2 expression in T47D *shmdm2* as compared to controls, correlated with reduced 53BP1 and MDC1 on chromatin by examining the chromatin bound 53BP1 and MDC1 (Fig. 1B), as well as the cellular localization levels of phospho-Ser25 53BP1 and phospho-Ser1778 53BP1 by immunofluorescence (Figs. 1C and 1D). Consistent with the SILAC data we found that in the MDM2-depleted T47D cell line there were reduced levels of both 53BP1 and MDC1 associated with chromatin (Figs. 1B and 1C). In addition, immunofluorescence revealed fewer phospho-53BP1 foci in both MDM2-depleted and MDMX-depleted cells compared to the vector control cells, a result consistent with the chromatin binding activity data as phosphorylation of 53BP1 happens when it is bound to chromatin. Thus, we confirmed our SILAC analysis which revealed a role for MDM2 for increasing 53BP1 recruitment to chromatin in cancer cells.

### MDM2 interacts with 53BP1 in vivo

The ability of MDM2 to interact with 53BP1 *in vivo* was examined using the Proximity Ligation Assay (PLA) within parental control, MDM2-depleted, and MDMX-depleted derivatives of two breast cancer cell lines: T47D and MDA-MB-231 (mtp53 R280K). We compared the expression levels of mtp53, 53BP1, and MDM2 within each cell line (Fig 2A), observing that T47D cells expressed ∼3-fold higher MDM2 levels due to the presence of the *MDM2* gene *SNP309 G/G* allele [48]. Because 53BP1 has been shown to co-immunoprecipitate with p53, we tested the ability of mtp53 in our T47D and MDA-MB-231 cell lines to form a PLA interaction with 53BP1 (Fig. 2B). We observed a mtp53-53BP1 PLA-interaction in both T47D and MDA-MB-231 control cells that was independent of MDM2 and MDMX, since in both MDMX-depleted and MDM2-depleted cells we did not observe a decrease in 53BP1-mtp53 PLA foci per nuclei. Importantly, we observed a mtp53-MDM2 PLA-interaction in vector control cells (Fig. 2C) that, in contrast to the mtp53-53BP1 PLA-interaction, was reduced in both the MDMX-depleted and MDM2-depleted cells. Lastly, we detected in both mtp53 breast cancer cell lines a significant MDM2-53BP1 PLA-interaction (Fig. 2D) that was neither diminished or enhanced in the MDMX-depleted cells, suggesting both forms of MDM2 (homodimer and heterodimer) are competent in this assay.

**Figure 2:**
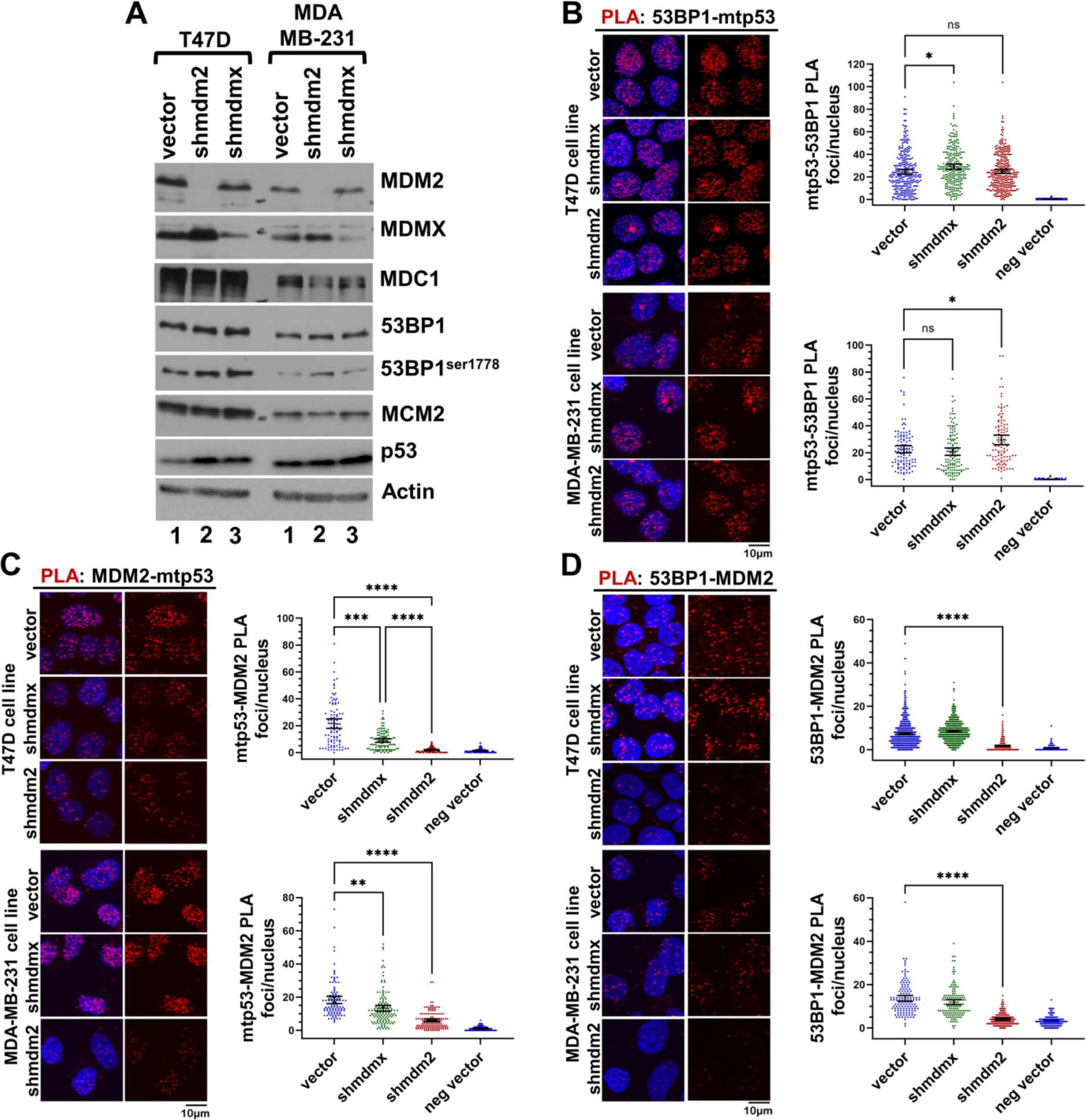
53BP1 in breast cancer cells interacts with both mtp53 and MDM2. **(A)** Abundance of mtp53, MDM2, MDMX, 53BP1, and MDC1 within extracts (20µg) prepared from T47D (mtp53 L194F) and MDA-MB-231 (mtp53 R280K) cell lines was determined by SDS-PAGE/western blot analysis. **(B-D)** Association of mtp53, MDM2, and 53BP1 *in vivo* measured using the Proximity Ligation Assay. PLA analysis of mtp53-53BP1 (panel B), MDM2-mtp53 (panel C), and MDM2-53BP1 (panel D) within T47D and MDA-MB-231 cells were performed as described in the materials and methods; primary antibodies are PLA rabbit anti-p53, PLA goat anti-53BP1, and mouse anti-MDM2 4B2. Confocal images for 4-6 fields for each were acquired and the number of PLA foci/nucleus from each cell population was determined using Cell Profiler. Graphs and statistical analyses (Kruskal-Wallis) of data from >100 cells in panel B, and >250 cells in panels C and D were prepared using Prism. Data representations are mean foci/nucleus with 95% confidence interval (n=3 for T47D cell line analyses, n=2 for MDA-MB-231 cell line analyses); p < .05 (*), p < .01 (**), p < .001 (***), p < .0001 (****), p > .05 (ns).

### The C-terminus of mtp53 is required for MDM2 interaction with 53BP1

To address the possibility that mtp53 helps to coordinate the close proximity for the identified MDM2-53BP1 foci, we analyzed the MDM2-53BP1 PLA-interaction within the breast cancer cell line MDA-MB-468 (expressing mtp53 R273H) and an MDA-MB-468 derivative in which CRISPR-Cas9 was used to delete the mtp53 MDM2-interacting C-terminal domain (Fig. 3A) [39]. The resulting mtp53 protein, R273H*fs*347Δ360-393 (referred to here as R273HΔC) lacks the C-terminal 10 amino acids of the oligomerization domain and the entire C-terminal regulatory domain, which includes a functional MDM2 binding site [49]. We reasoned that we could compare the MDM2-53BP1 PLA interaction efficiency between the cell lines to assess the impact of loss of MDM2 interacting with mtp53 on the interaction of MDM2 with 53BP1. Western blot analysis of extracts from each cell line showed comparable levels of 53BP1 in the two cell lines but in the CRISPR-Cas9 derivative we found 3-fold lower mtp53 R273HΔC. We then asked if any mtp53 R273HΔC was detected in a complex with MDM2 by co-immunoprecipitation with two different MDM2 antibodies (Fig. 3C). We found that both MDM2 antibodies pulled down full length mtp53 R273H from the parental cell line extract, we were unable to detect co-IP of mtp53 R273HΔC from the CRISPR derivative cell line extract (Fig. 3C, compare lane 4 to lane 6 and lane 7 with twice the amount of extract used and lane 11 to lane 13 and lane 14 with twice the amount of extract used). This indicated that loss of the C-terminus of mtp53 R273H prevented stable binding of mtp53 to MDM2. However, while we could detect a stable interaction in whole cell extracts between MDM2 and mtp53 we were only able to detect a similar interaction between MDM2 and 53BP1 by PLA analysis. This suggested that mtp53 protein might regulate the interaction. When using PLA, we detected the interaction of mtp53 with 53BP1 in both the MDA-MB-468 cell line and the mtp53 R273HΔC derivative loss of the C-terminus with a 2.5-fold reduction that was consistent with the reduced mtp53 expression (Fig, 3D). When the mtp53-MDM2 PLA interaction was measured we detected a 5.1-fold reduction in the mean PLA foci per nucleus in the R273HΔC-expressing cell line compared to the mtp53 R273H-expressing parental control (8.8 PLA foci per nucleus in cells with mtp53 R273HΔC versus 44.9 in cells with mtp53 R273H) (Fig. 3E). This showed a residual interaction that was not seen with the co-IP data but still indicated compromised binding of mtp53 R273HΔC to MDM2. The comparative MDM2-53BP1 PLA interaction for the two cell lines revealed a 2.7-fold decrease in the mean PLA foci per nucleus in the mtp53 R273HΔC-expressing cell line compared to the mtp53 R273H-expressing parental control (5.7 for mtp53 R273HΔC versus 15.3 for mtp53 R273H). This decrease was consistent with the reduction in the mtp53-MDM2 PLA-interaction when normalized for the expression levels of each mtp53. We therefore further evaluated if the mtp53-53BP1 interaction would target MDM2 to 53BP1. We addressed this question further by using the small molecule p53-MDM2 disrupter Nutlin 3a, which is known to allow release and activation of wtp53 [49, 50]

**Figure 3:**
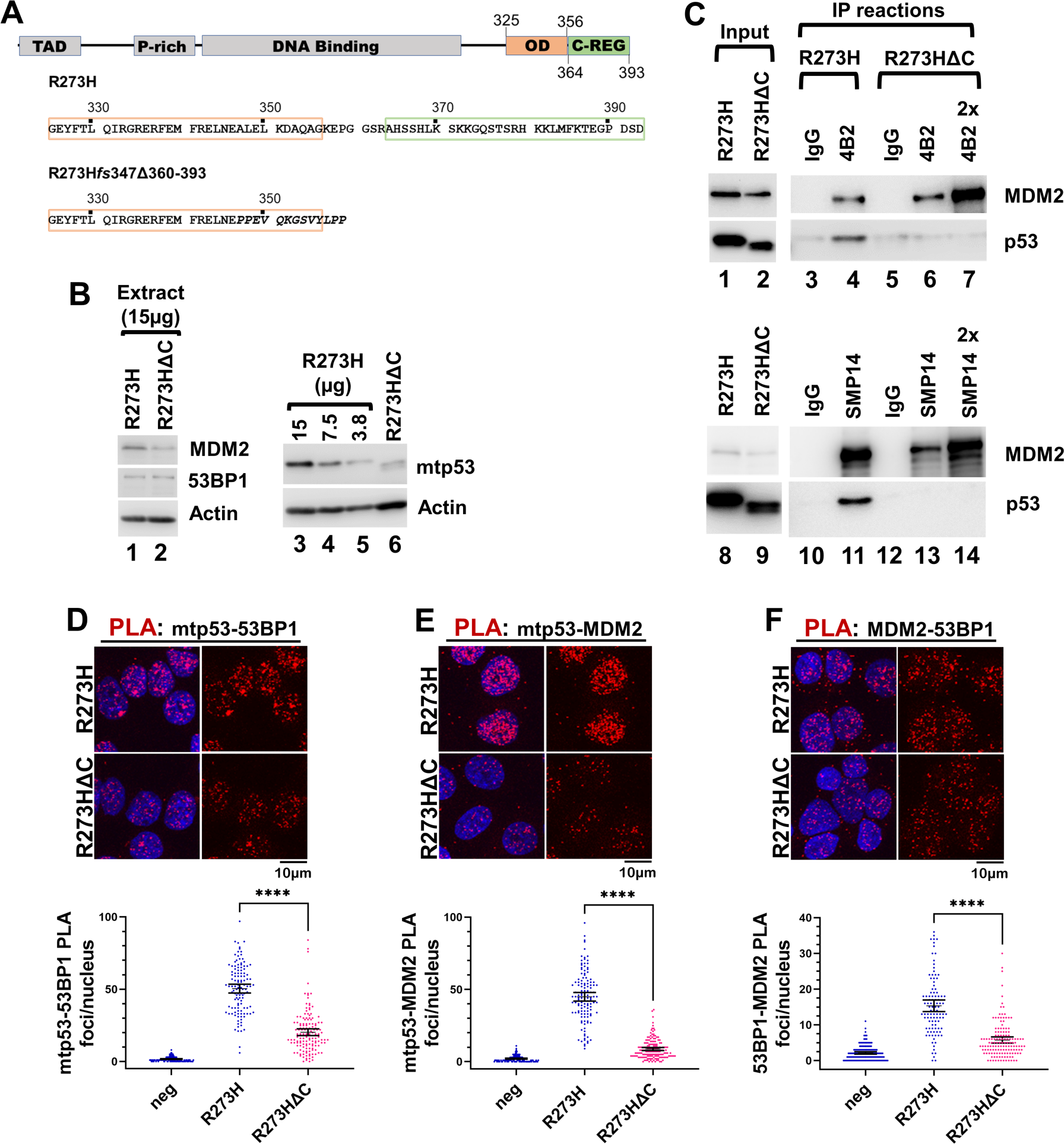
The MDM2-53BP1 interaction is promoted by the mtp53 C-terminus. **(A)** The C-terminus of mtp53 R273H within MDA-MB-468 was modified using CRISPR/Cas9 to create the cell line MDA-MB-468 R273H*fs*347Δ360-393 (termed G6; mtp53 derivative R273HΔC) [39]. **(B)** Relative protein levels of MDM2, 53BP1, and mtp53 within MDA-MB-468 (15 µg, 7.5 µg, and 3.75 µg (labeled 3.8) and MDA-MB-468 R273H*fs*347Δ360-393 (15µg) cell lines was examined by SDS-PAGE/western blot analysis. **(C)** Loss of mtp53 C-terminus disrupts mtp53 co-immunoprecipitation (IP) with MDM2. Extracts from MDA-MB-468 and MDA-MB-468 R273H*fs*347Δ360-393 cell lines were incubated with either normal mouse IgG (negative control) or anti-MDM2 antibodies 4B2 (upper panel) or SMP14 (lower panel), and the resulting IP reactions were examined for the presence of MDM2, and mtp53 by SDS-PAGE/western blot analysis. In the upper panel, reactions were performed using 800 µg of each extract, and gel lanes contain 10% of each immunoprecipitate and 2% of the IP reaction input (due to the vast excess of mtp53 compared to MDM2 within MDA-MB-468 a lighter exposure of mtp53 input is presented). In the lower panel, reactions were performed using 1600 µg of each extract, and gel lanes represent 12.5% of each immunoprecipitate and 0.5% of the IP reaction input. In both panels, an extra lane containing twice the amount of the MDA-MB-468 R273H*fs*347Δ360-393 extract MDM2 IP was loaded (labeled 2x; 20% of the MDA-MB-468 R273H*fs*347Δ360-393 4B2 IP and 25% of G6 SMP14 IP). **(D-F)** PLA analysis of mtp53-53BP1 (panel D), MDM2-mtp53 (panel E), and MDM2-53BP1 (Panel F) in MDA-MB-468 and MDA-MB-468 R273H*fs*347Δ360-393. Close proximity of the indicated proteins was measured using the proximity ligation assay performed as described in the materials and methods; primary antibodies used are PLA rabbit anti-p53, PLA goat anti-53BP1, and mouse anti-MDM2 2A9. Confocal images for 3-5 fields for each were acquired and the number of PLA foci/nucleus from each cell population was determined using Cell Profiler. Graphs and statistical analyses (Kruskal-Wallis) of data from 100-150 cells in each panel was prepared using Prism. Data representations are mean foci/nucleus with 95% confidence interval (n=2); p < .05 (*), p < .01 (**), p < .001 (***), p < .0001 (****), p > .05 (ns).

### Both the MDM2-mtp53 and MDM2-53BP1 PLA-interactions are Nutlin 3a sensitive

The comparison experiments using the MDA-MB-468 expressing mtp53 R273HΔC versus full-length mtp53 R273H cell lines suggested mtp53 helped to direct MDM2 into close proximity with 53BP1. We therefore asked if Nutlin 3a treatment of T47D cells could phenocopy the reduced MDM2-53BP1 PLA observation (Fig. 4). We first asked if the cell cycle of T47D cells expressing mtp53 was perturbed by 10µM Nutlin 3a treatment by comparing treated T47D cells to Nutlin 3a treated MCF7 cells (expressing wtp53). In wtp53 expressing cells, Nutlin 3a induces cell cycle checkpoints and apoptosis by increasing p53 by blocking the MDM2-mediated degradation, and consequently up-regulation of the p53 protein and transactivation activity (Supplementary S1 (Panels A and B). We treated MCF7 and all three variant T47D cell lines with 10µM Nutlin 3a, and then examined each cell population by western blotting for levels of 53BP1, MDM2, MDMX, p53, cyclin A, and cyclin B (Fig. 4A), and EdU labeling and immunofluorescence for Cyclin A and flow cytometry (Fig. 4B and Supplementary S1 Panel C). The Nutlin 3a-treated MCF7 extracts in western blot analysis revealed a significant increase in the level of p53 and induction of MDM2 protein indicative of wtp53 activation (Fig. 4A compare lane 1 to lanes 2 and 3). Upon Nutlin 3a treatment all three T47D cells showed no such p53 activation of the p53 signal transduction pathway (Fig. 4A lanes 4 through 9). Further confirmation of the bio-activity of the Nutlin 3a was observed when we examined the levels of the S-G2 cell cycle markers Cyclin A and Cyclin B when compared to the extract from untreated MCF7 cells (Fig. 4A lanes 1 to 3), and a reduction in the levels of Cyclin A and B were observed in the extract from 24hr Nutlin 3a-treated cells (Fig. 4A lane 3). This was consistent with a decrease in the S-G2 cell cycle fraction and a G1/S arrest. In contrast, analysis of extracts from T47D cell populations revealed comparable levels of Cyclin A and Cyclin B within Nutlin 3a-treated extracts (Fig.4A lanes 4 through 9), suggesting no significant change in the S-G2 cell cycle fraction. We also examined Nutlin 3a treated T47D cells by immunofluorescence microscopy for a G1/S arrest by measuring the percentage of cells that incorporated EdU and expressed Cyclin A (Fig. 4B). All three Nutlin 3a treated T47D cell lines exhibited no inhibition of EdU incorporation or significant change in the percentage of S-G2 cells using this method or by flow cytometry (Fig. 4B and Supplementary S1 Panel C). We thus concluded that Nutlin 3a treatment of mtp53-expressing T47D cells did not induce a measurable cell cycle arrest phenotype.

**Figure 4:**
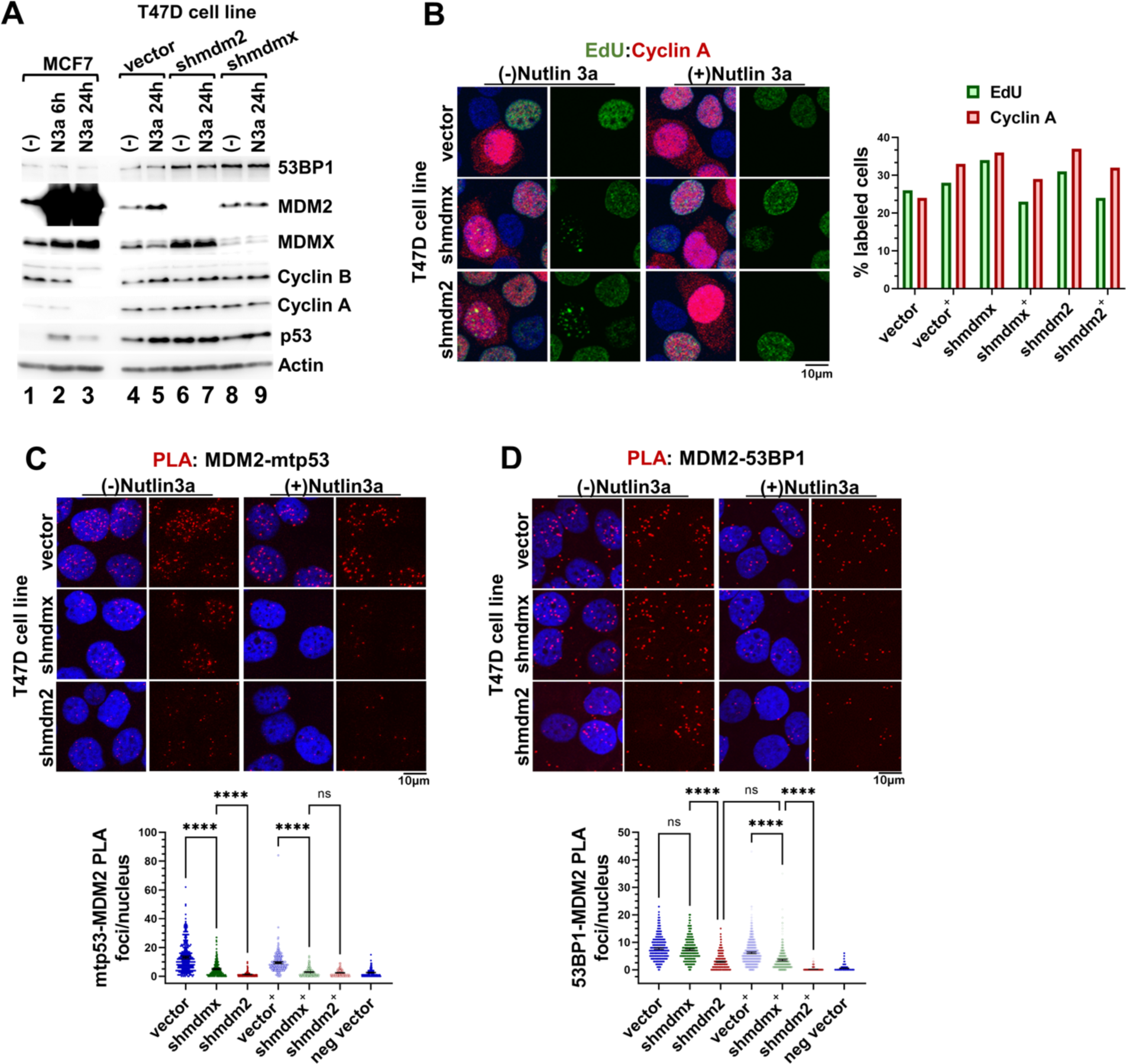
The MDM2-mtp53 and MDM2-53BP1 PLA-interactions are Nutlin 3a sensitive. **(A)** Abundance of mtp53, MDM2, MDMX, 53BP1, Cyclin A, and Cyclin B within extracts from MCF7 and T47D cell line populations treated with Nutlin 3a (N3a) for 24 hr and untreated populations were compared by western blot analysis. **(B)** Treatment of T47D cell lines with Nultin 3a does not inhibit cell cycle progression. T47D cell line populations untreated (-) and treated (+) with 10µM Nutlin 3a for 24hr were incubated with EdU for 15 min, and the percentage of S-G2 phase cells for each population (250-285 cells counted) was determined by confocal microscopy after (1) labeling incorporated EdU with Alexa Fluor 647 (pseudo-colored green) and (2) immunofluorescence staining for Cyclin A (mouse anti-Cyclin A detected with anti-mouse Alexa Fluor 594). **(C and D)** Disruption of mtp53-MDM2 and 53BP1-MDM2 PLA interactions in Nutlin 3a-treated T47D cells. MDM2-mtp53 (Panel C) and MDM2-53BP1 (Panel D) interactions within T47D cell lines untreated or post 24hr treatment with 10µM Nutlin3a were measured using the proximity ligation assay performed as described in the materials and methods; primary antibodies used are PLA rabbit anti-p53, PLA goat anti-53BP1, and mouse anti-MDM2 4B2. Confocal images for 4-6 fields for each were acquired and the number of PLA foci/nucleus from each cell population was determined using Cell Profiler. Graphs and statistical analyses (Kruskal-Wallis) of data from >100 cells in Panel C and >300 cells in Panel D prepared using Prism. Data representations are mean foci/nucleus with 95% confidence interval (n=2 for panel C and n=3 for panel D); p < .05 (*), p < .01 (**), p < .001 (***), p < .0001 (****), p > .05 (ns).

We then asked if treating the T47D cells with Nutlin 3a, reduced the PLA foci for MDM2-mtp53 (Fig. 4C) and MDM2-53BP1 (Fig. 4 D). We observed that before Nutlin 3a treatment the T47D MDM2-mtp53 PLA-interaction was decreased in the MDMX-depleted cells and was significantly more reduced in the MDM2-depleted cells (Fig. 4C). However, following treatment with Nutlin 3a we observed an equally significant reduction in the MDM2-mtp53 PLA-interaction in both the cells with MDMX or MDM2 depletion (Fig. 4C). Strikingly, in the absence of MDMX we observed that Nutlin 3a treatment completely blocked the MDM2-p53 PLA foci (Fig. 4C).

We next measured the MDM2-53BP1 PLA-interaction before and after 24hr of treatment with Nutlin 3a (Fig. 4D). We found a comparable number of MDM2-53BP1 PLA foci per nucleus for the cells with and without and MDMX depletion but saw that the MDM2-depleted cells had reduced MDM2-53BP1 PLA foci per nucleus (Fig. 4D). However, within the Nutlin 3a-treated MDMX-depleted cells compared to Nutlin 3a treated control cells, we observed a significant decrease in the MDM2-53BP1 PLA-interaction, and for the MDM2-depleted cells we observed a further decrease (Fig.4D). Taken together, this analysis suggests that the mtp53-MDM2 complex, stabilized by MDMX, targets MDM2 to be in close proximity with 53BP1 on chromatin.

### MDM2 regulates the interaction between 53BP1 and MDC1

Given that our SILAC screen also showed a reduction in the DNA repair protein MDC1 (Figs.1A and 1B), and was performed using chromatin from cells not exposed to exogenous DNA damage agents, we posited that MDM2-influenced increased recruitment of MDC1 and 53BP1 that might be a persistent coordinated replication stress alleviator. Using the PLA assay within the T47D cell lines we measured the MDC1-53BP1 PLA foci per nucleus interaction in the absence, and presence, of the mtp53-MDM2 disruptor or in the presence of the DNA damaging agent etoposide (Fig. 5).

**Figure 5:**
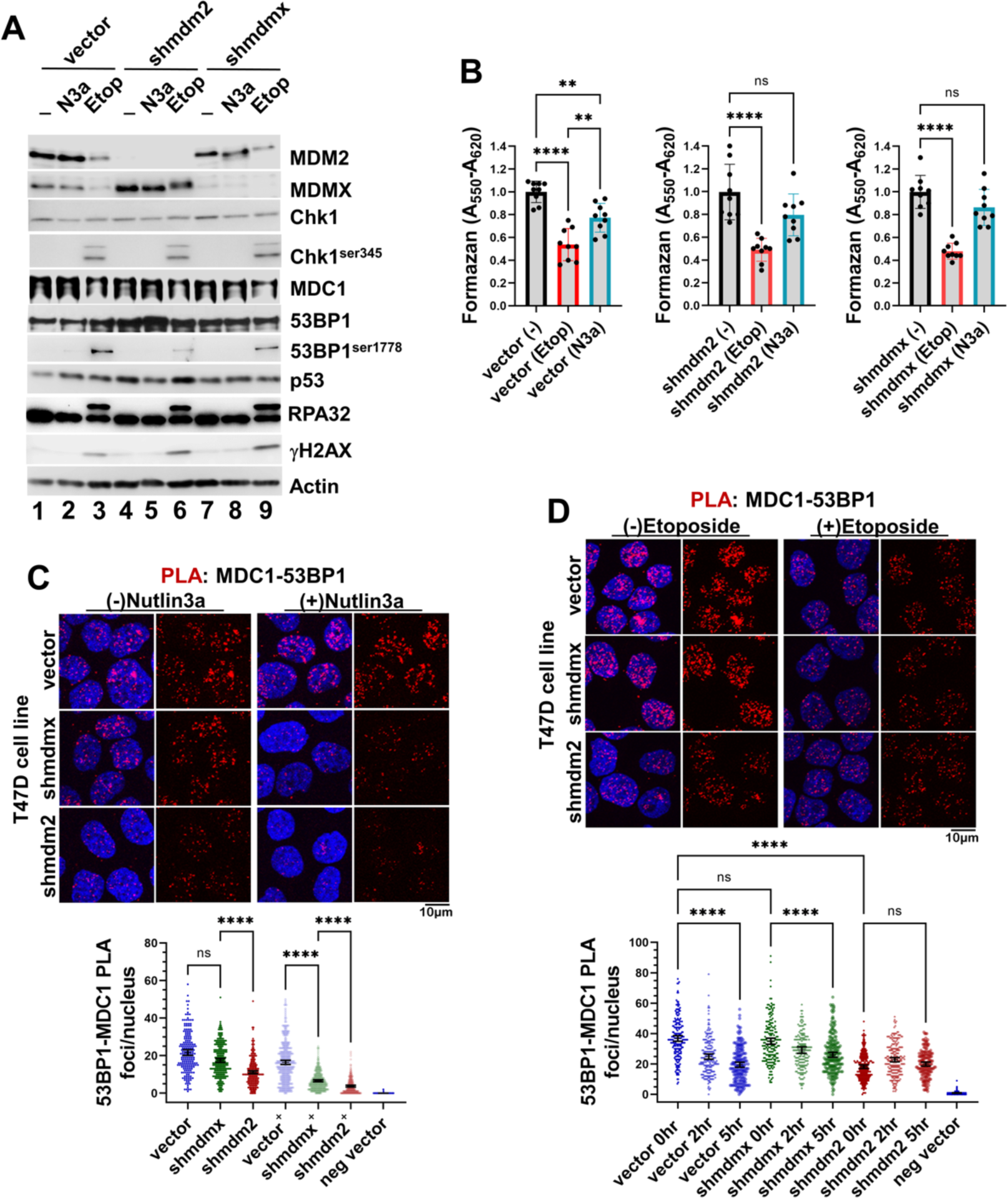
The MDM2-53BP1 PLA interactions are diminished by loss of MDM2 function. Activation of the DNA damage response in T47D cell lines by Etoposide but not Nutlin-3a. **(A)** Extracts (20µg) from T47D cell lines treated with either Etoposide for 5hr (Etop; 50µM), or Nutlin 3a for 24hr (N3a; 10µM) were analyzed for the indicated proteins by western blotting. **(B)** Cell viability of T47D cell lines treated with either 50µM Etoposide or 10µM Nutlin 3a for 24hr was measured using the MTT assay. **(C and D)** MDC1-53BP1 *in vivo* interactions were measured within each T47D cell line 24hr-post Nutlin 3a treatment **(Panel C)** or the indicated time points post Etoposide treatment **(Panel D)** using the proximity ligation assay as described in the materials and methods; primary antibodies used are PLA rabbit anti-p53, PLA goat anti-53BP1, PLA rabbit anti-MDC1, and mouse anti-MDM2 4B2. Confocal images for 3-6 fields for each were acquired and the number of PLA foci/nucleus from each cell population was determined using Cell Profiler. Graphs and statistical analyses (Kruskal-Wallis) of data from >250 cells in Panel C and >200 cells in Panel D prepared using Prism. Data representations are mean foci/nucleus with 95% confidence interval (n=3 for panel C and n=2 for panel D); p < .05 (*), p < .01 (**), p < .001 (***), p < .0001 (****), p > .05 (ns).

We analyzed for DNA damage checkpoint activation by western blotting extracts prepared from each T47D cell line either untreated or treated for 24 hr-post 10µM Nutlin 3a or 5 hr-post 50µM etoposide, for phospho-Ser1778 53BP1, γH2AX, phospho-Chk1, as well as reduction in the levels of MDM2 and MDMX (Fig. 5A). Only the etoposide treated cells demonstrated DNA damage response pathway activation of phospho-Chk1, phospho-Ser1778 53BP1, and γH2AX (Fig. 5A, see lanes 3, 6, and 9). The etoposide-induced DNA damage also resulted in reduction in both MDM2 and MDMX (Fig. 5A, lanes 3 and 9), with loss of the MDMX protein dependent on MDM2 function (Fig. 5A, lane 6). Levels of phospho-Chk1, phospho-Ser1778 53BP1, and γH2AX from Nutlin 3a treated cells were found comparable to the untreated T47D cell lines (Fig. 5A, lanes 2, 5, and 8). Moreover, the MTT assay revealed all three T47D cell lines displayed comparable sensitivity to etoposide, and little to no sensitivity to Nutlin 3a (Fig. 5B).

When close proximity of MDC1 and 53BP1 was assessed in our cell lines, the vector control cells demonstrated clear MDC1-53BP1 PLA nuclear foci (Fig. 5C). Compared to the T47D control cells the MDM2-depleted cells, but not the MDMX-depleted cells, displayed a significant reduction in the number of MDC1-53BP1 PLA foci per nucleus (Fig. 5C, see dark green versus dark red points). However, exposure to Nutlin 3a caused the MDMX-depleted cell line to show a striking reduction in the number of MDC1-53BP1 PLA foci per nucleus (Fig. 5C, see light green and light red points). These results mirrored those obtained in the Nutlin 3a analysis of the PLA-interaction for MDM2-53BP1, and provides further support for a mtp53-MDM2-MDMX circuit driving 53BP1 to modulate its activity on chromatin. Because the observed down-regulation of the MDC1-53BP1 PLA-interaction was measured in the absence of DNA damage response activation, we also examined the MDC1-53BP1 PLA-interaction after etoposide-induced DNA damage (Fig. 5D). Etoposide treatment creates the chromatin modification necessary for 53BP1 recruitment independent of MDC1. As such, in the vector control cells treated with etoposide we observed a time dependent down-regulation of MDC1-53BP1 PLA foci (Fig. 5D, see blue points for 2hr and 5hr etoposide treatment). In both the control and MDMX-depleted cells 5hr post etoposide DNA damage response activation, when significant reduction in MDM2 (and MDMX) levels were observed, the PLA foci levels approached levels measured in the untreated MDM2-depleted cells (Fig. 5D, see blue and green points for 2hr and 5hr etoposide treatment). In stark contrast, the MDC1-53BP1 PLA foci levels were so low within MDM2-depleted cells that after etoposide treatment they remained unchanged (Fig. 5D, see red points for 2hr and 5hr etoposide treatment). These data suggest that the loss of MDC1-53BP1 PLA foci was due to the loss in MDM2 protein. Taken together, these results suggest MDM2 functions to promote 53BP1 chromatin residency mediated through an interaction with MDC1 in the absence of DNA damage activation. Upon DNA damage pathway activation, PI3 kinase activity causes reduction of MDM2 and an increase in histone-PTMs that promote an alternative pathway for 53BP1 chromatin recruitment. There are numerous pathways that help facilitate repair of replicating DNA and mtp53 participates with poly-ADP ribose polymerase (PARP) in such a process [27]. We therefore asked if MDM2, in the presence of mtp53, might coordinate to regulate PARP activity. This would not be surprising because MDM2, in the context of wtp53, has been shown to influence PARP activity to enhance replication fork progression [32]

### MDM2 Protein also Regulated Chromatin PARylation

Although MDM2 has been reported to be a down-regulator of PARP1 by ubiquitin mediated proteolysis in the context of wtp53 [32], such a function has not been reported in the context of mtp53. We therefore asked if inhibition of the MDM2-p53 interaction as described in MCF7 cells induced decreased levels of PARP1. We first reproduced results of reduction of PARP 1 protein by treating MCF7 cells with Nutlin 3a (Supplementary S1 Panel B MCF7 cells 24 hr Nutlin 3a treatment). In contrast to MCF7 cells in which we observed reduction of PARP1, in Nutin3a-treated T47D control and MDM2 or MDMX knockdown cells, we observed that the PARP1 protein level remained unchanged (Supplementary S1 Panel B T47D cells 24 hr Nutlin 3a or Aphidicolin treatment). In MCF7 cells the interaction between wtp53 and 53BP1 as well as wtp53 and MDM2 remained intact (Supplementary S1 Panels D and E), indicating loss of PARP1 after 24 hr Nutlin 3a treatment did not perturb the p53-53BP1 interaction. We have shown that mtp53 interacts with PARP1 in breast cancer cells and that mtp53 expression increases cancer cell sensitivity to combination PARP inhibitor (PARPi) talazoparib with temozolomide treatment [27, 28, 51, 52]. In T47D cells we assessed the influence of depletion of either MDMX or MDM2 on cytosolic and chromatin PARP protein level and PARP enzymatic activity (Figs. 6A and 6B). The three different T47D cell lines were either left untreated, treated with the combination of the PARPi talazaparib in combination with the DNA damage agent temozolomide, or treated with the combination of the PARPi and temozolomide and then allowed 24 hr of DNA repair. In all three cell lines with the combination of a PARPi and temozolomide, or when treatment was followed by 24 hr of DNA repair, we observed a robust decrease in PARylation of proteins (Fig. 6A for cytosol see PAR panel to compare PARylated proteins in lanes 1 to 3, to reduced PARylated proteins in lanes 4 through 9; and 6B for chromatin see PAR panel to compare PARylated proteins in lanes 1 to 3, to reduced PARylated proteins in lanes 4 through 9). Importantly, in the untreated chromatin and cytoplasmic fractions we saw that the depletion of MDM2 resulted in increased PARylated protein (Figs. 6A and 6B, see PAR panels lane 3). Moreover when the PARylation of proteins was partially restored after 24 hr following removal of the drug, we observed that with time given for DNA repair resulted in increased chromatin PARylation (Fig 6B, lanes 7 through 9). The 24 hr without the drug cell extract preparations showed increased DNA stress markers of high Rad51, γH2AX, and RPA32 chromatin recruitment (Fig. 6B, compare lane 7, 8, and 9). The depletion of MDM2 caused decrease chromatin associated MDC1 before, during, and after PARPi treatments (Fig. 6B, lanes 3, 6, and 9). We also observed that depletion of MDM2 in MDA-MB-231 activated PARP enzymatic activity, and that following PARPi treatment the MDM2 depletion assisted in PARylation reactivation (data not shown). Both the 53BP1 and the PARP repair pathways have been shown to participate in the maturation of Okazaki fragments during DNA replication [53, 54]. Taken together these results suggest that MDM2 works together with mtp53 to assist in coordinating the responses initiated in cancer cells during DNA replication stress. We posit that this coordination is responsible for the cancer cell CPR response to survive accumulated DNA damage that would cause most cells to die.

**Figure 6:**
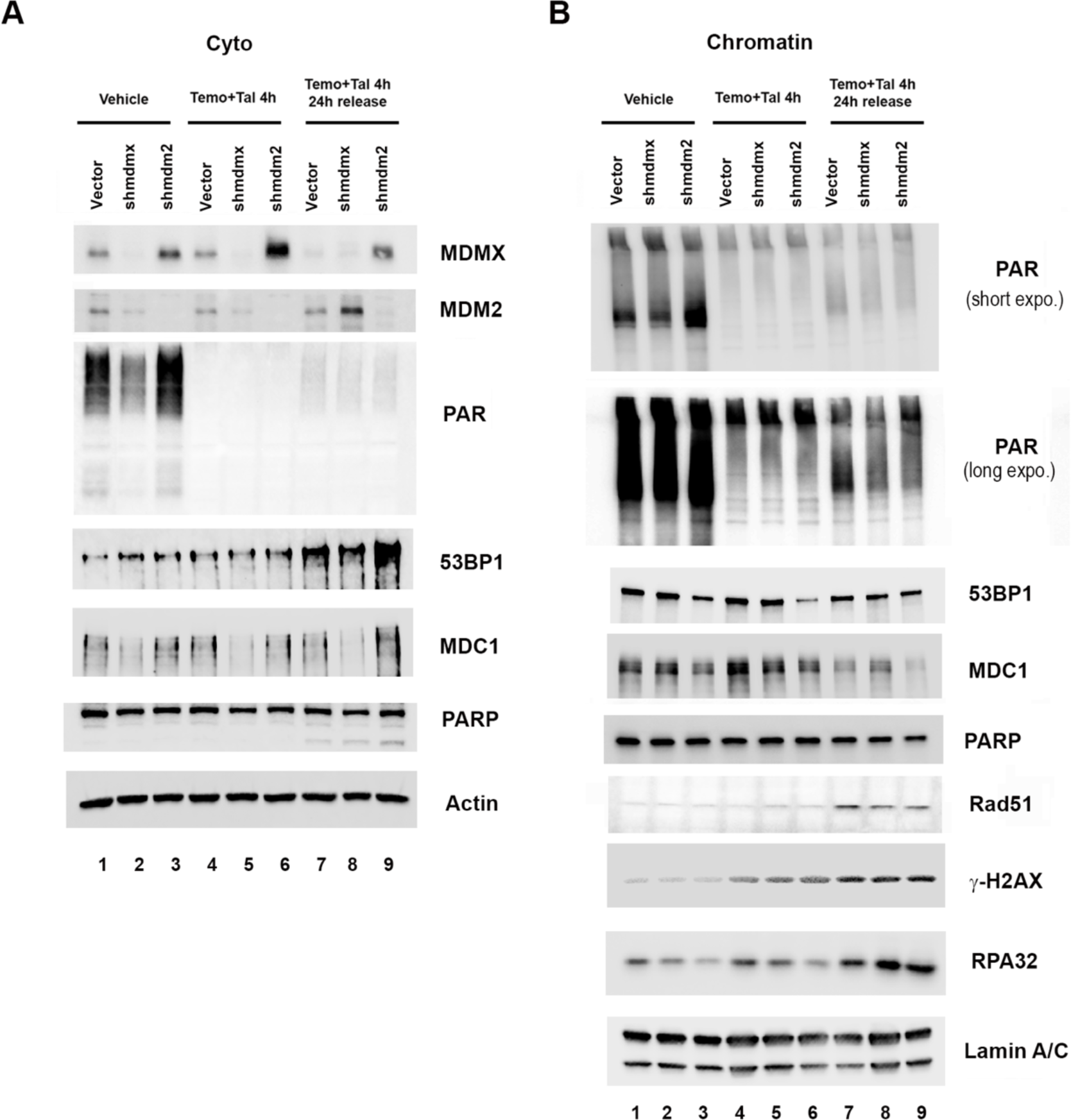
MDM2 Reduces Poly(ADP-Ribose) (PAR) Levels of Chromatin Bound Proteins. Cytosolic (A) and Chromatin (B) fractions were prepared from T47D cells with constitutive *shmdm2*, *shmdmx*, or mir30 shRNA-expressing vector cells treated with either vehicle (DMSO), or a combination of 1mM temozolomide plus10 μM talazoparib (Temo+Tal) for 4 h, or combination of 1mM temozolomide plus10 μM talazoparib (Temo+Tal) for 4 h then replace with fresh media for additional 24 h. 10 μg of cytosolic or chromatin protein was loaded on a SDS-PAGE and protein levels were determined by Western blot analysis using the indicated antibodies. Abbreviations: DNA double strand breaks (DSBs), Immunofluorescence microscopy (IF), NHEJ, HR, MDC1, MDM2, MTT, etc.

## Discussion

Herein we show that MDM2 works together with mtp53 (in a number of different breast cancer cell lines) to coordinate the response of normal replication stress using a 53BP1-MDC1 pathway. It is known that MDM2 interacts with MDC1 and the Mre11-Rad50-Nbs1 complex (MRN) DNA repair machinery [12, 47, 55]. However, before this work, no known connection had been described between MDM2 and 53BP1. Using a SILAC screen we determined that MDM2 was required for chromatin recruitment of phosphorylated MDC1 and 53BP1 (see Fig. 1). Furthermore, we observed a clear mtp53-dependent interaction of MDM2 with 53BP1 (see Figs. 2, 3, and 4).

When double-strand DNA breaks are generated, the damage is repaired by either the non-homologous end joining (NHEJ) or the homologous recombination (HR) pathway, with 53BP1 blocking HR and promoting NHEJ [56]. The interaction between 53BP1 and MDC1 is required for recruitment of 53BP1 to DNA breaks [57]. Rad51 directs HR which can only occur during S or G2 cell cycle stages, when cells have generated sister chromatids [58]. In G1 and G2, the 53BP1 protein drives predominately NHEJ [58]. However, 53BP1 also participates in reducing S-phase DNA damage and mitigating replication stress [59]. A maximal DNA damage response function requires MDC1-mediated recruitment of 53BP1 to chromatin through phospho-Ser139 H2AX [34]. However, when MDC1 interacts with chromatin in the absence of DNA damage, it does so in a γH2AX-independent manner through a Proline-Serine-Threonine repeat (PST) domain that contributes to 53BP1 chromatin retention [34, 35]. We carried out experiments in the absence, versus the addition, of exogenous DNA damaging agents and found an MDM2-dependent interaction of 53BP1 interacting with MDC1 (Fig. 5). Damaged DNA creates chromatin territories by MDC1 alternatively regulating the DNA damage sensor MRN and Rad51 to promote HR, or 53BP1 to interact with RIF1 and the Shieldin complex to protect DNA ends from nucleolytic attack by MRN and subsequently promote NHEJ repair [56].

MDM2 is known to negatively regulate wtp53 function, however in the context of mtp53 the data presented here suggest a role reversal, in which mtp53 regulates MDM2. Using the p53-MDM2 interaction disruptor Nutlin 3a, we phenocopied the loss of the 53BP1-MDC1 PLA interaction displayed in MDM2-depleted T47D cells (Fig. 5). This suggests the MDM2-mtp53 complex mediates the 53BP1-MDM2 interaction. Additionally, we provide evidence for MDM2-mediated negative regulation of PARP1 activity in the context of mtp53, as loss of MDM2 in T47D and MDA-MB-231 cells resulted in enhanced cellular protein PARylation (Fig. 6 and data not shown). Interestingly, in this context the PARP1 protein level was stable as it was Nutlin 3a treated with T47D cell. In contrast the MCF7 cells treated with Nutlin 3a underwent MDM2-promoted ubiquitin mediated proteolysis of PARP ([32] and Supplementary Fig. S1 Panel B). Thus, regulation of MDM2 activity may augment GOF mtp53 properties. Indeed, we speculate that the mtp53-MDM2 complex is the active form of mtp53 that mediates DNA replication, and DNA repair, and mtp53 GOF activities. The MDC1-53BP1 interaction promoted by MDM2 with mtp53 could be viewed as a mtp53 GOF activity that bypasses the need for PI3 kinase (ATM/ATR) activation. This interpretation is consistent with that proposed for the mtp53-TopBP1 interaction that drives DNA replication [30]. In this mtp53 GOF pathway, mtp53 abrogates the requirement for both PI3 kinase and Cyclin/Cdk activity, which promotes the TopBP1-Treslin interaction and replisome assembly [30]. Recently, the active form of Treslin in DNA replication has been shown to be a complex with MTBP, a protein identified as an MDM2 binding protein. In light of this discovery, a role for MDM2 in the mtp53-TopBP1-Treslin pathway merits investigation.

Mutant p53 participates in cancer cell DNA replication, promotes tumorigenesis, and activates metastasis [27, 36, 60, 61]. Our previous work has focused on the interaction of mtp53 with PARP and PARylated chromatin [27, 28, 51, 52]. MDM2 is known for regulating DNA replication, chromatin, and DNA repair by down-regulating PARP and wtp53 [32, 62]. However the ability of mtp53 to work with MDM2 to influence DNA replication and repair is addressed herein for the first time. As such, upon discovering the ability of MDM2 to recruit MDC1 and 53BP1 to chromatin we became particularly interested in how MDM2 regulated mtp53-based PARP1 functionality. In conditions of cancer cells with mtp53 (using two different breast cancer cell lines) we found that reduction of MDM2 did not increase PARP protein level but did result in more PARylation of chromatin (Fig. 6 and data not shown). This suggests that in the absence of MDM2, when MDC1 and 53BP1 levels on chromatin are decreased, the PARP DNA repair pathway is activated. This contrasts with what is seen in cells with wtp53, which is that the inhibition of MDM2 interacting with wtp53 causes both an increase in wtp53 and MDM2 protein, and a decrease in total PARP levels because the MDM2 then works on PARP as an E3 ubiquitin ligase (see Supplementary Fig. S1 Panel B). The scenario in cancer cells for mtp53 and PARP is that the depletion of MDM2 does not cause either mtp53 or PARP levels to dramatically increase (see Figs. 1 through 6 and Supplementary S1). This may be because GOF mtp53 protects cells from MDM2-mediated PARP degradation. However, the MDM2 interaction with PARP appear to function to reduce PARP enzymatic activity. Future experiments are required to elucidate the cross-talk between mtp53, 53BP1, MDC1 and PARP in the context of cancer cell DNA replication stress. 53BP1 is a critical mediator of the wtp53-dependent cytotoxic effect of Nutlin 3a but in the presence of mtp53 no such 53BP1 cytotoxic influence occurs. This suggests that the ability to switch between a 53BP1 pathway and the PARP DNA damage pathway in Nutlin 3a treated mtp53 expressing cells may provide a cell survival advantage. Together mtp53, 53BP1, MDC1 and PARP proteins may coordinate in cancer cells to facilitate CPR to allow them to survive with replication stress and chromosomal instability that then causes an immunosuppressive tumor microenvironment [36].

## Acknowledgements

This work was supported by The Breast Cancer Research Foundation BCRF-20–011, BCRF-21–011, BCRF-22-011, and BCRF-23-011 to J. Bargonetti, and by National Cancer Institute of the National Institutes of Health under Award Number R01CA239603 to J. Bargonetti. This work was also partially supported by the TUFCCC/HC Regional Comprehensive Cancer Health Disparity Partnership from the NIH U54 CA221704. The content is solely the responsibility of the authors and does not necessarily represent the official views of the National Institutes of Health. We thank Katherine Harmon for creating visual models using BioRender and for proof reading the manuscript, and also thank students Emily Chan, Shuhong Jiang, Jessica Das, Amanda Larracuente, Khalikuz Mannan and members of the Bargonetti team for their efforts in support of this research.

## Notes

http://borreliabase.org/~wgqiu/clickme-khalikuz/temp-Points.html

### Competing Interest Statement

The authors have declared no competing interest.

