## Supplementary S1 for "A CANCER PERSISTENT DNA REPAIR CIRCUIT DRIVEN BY MDM2, MDM4 (MDMX), AND MUTANT P53 FOR RECRUITMENT OF MDC1 AND 53BP1 TO CHROMATIN"

### Supplementary Figure 1 (S1) Panels A-F

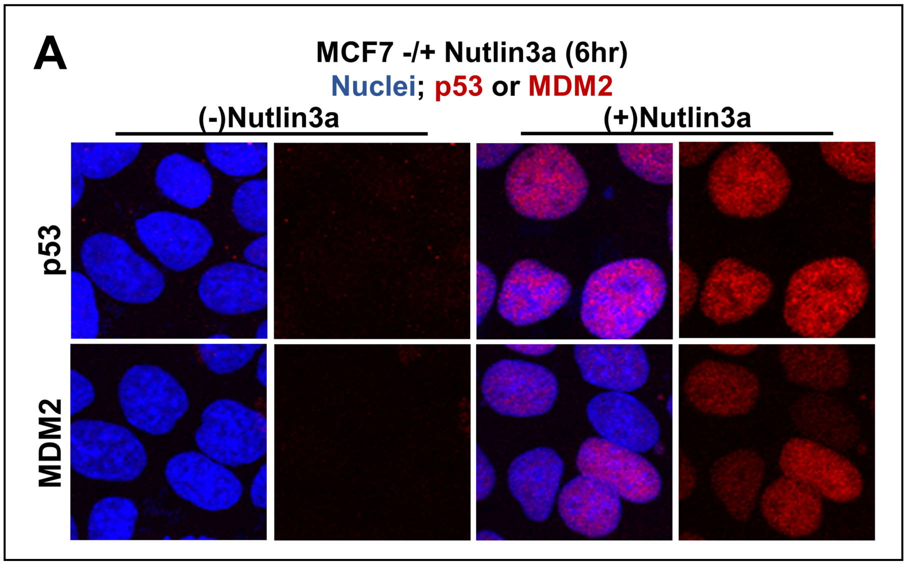


**Panel A:** *Nutlin 3a induction of wt p53 stabilization and transactivation in MCF7 cells.*

Immunofluorescence analysis for protein levels of p53 (detected with DO1) and MDM2 (detected with 4B2) within MCF7 cell cultures (50-60% confluency) untreated or treated with 10μM Nutlin 3a for 6hrs. Primary antibodies were detected with anti-mouse Alexa Fluor 594 (red) and nuclei were stained with Hoescht 33342 (blue).

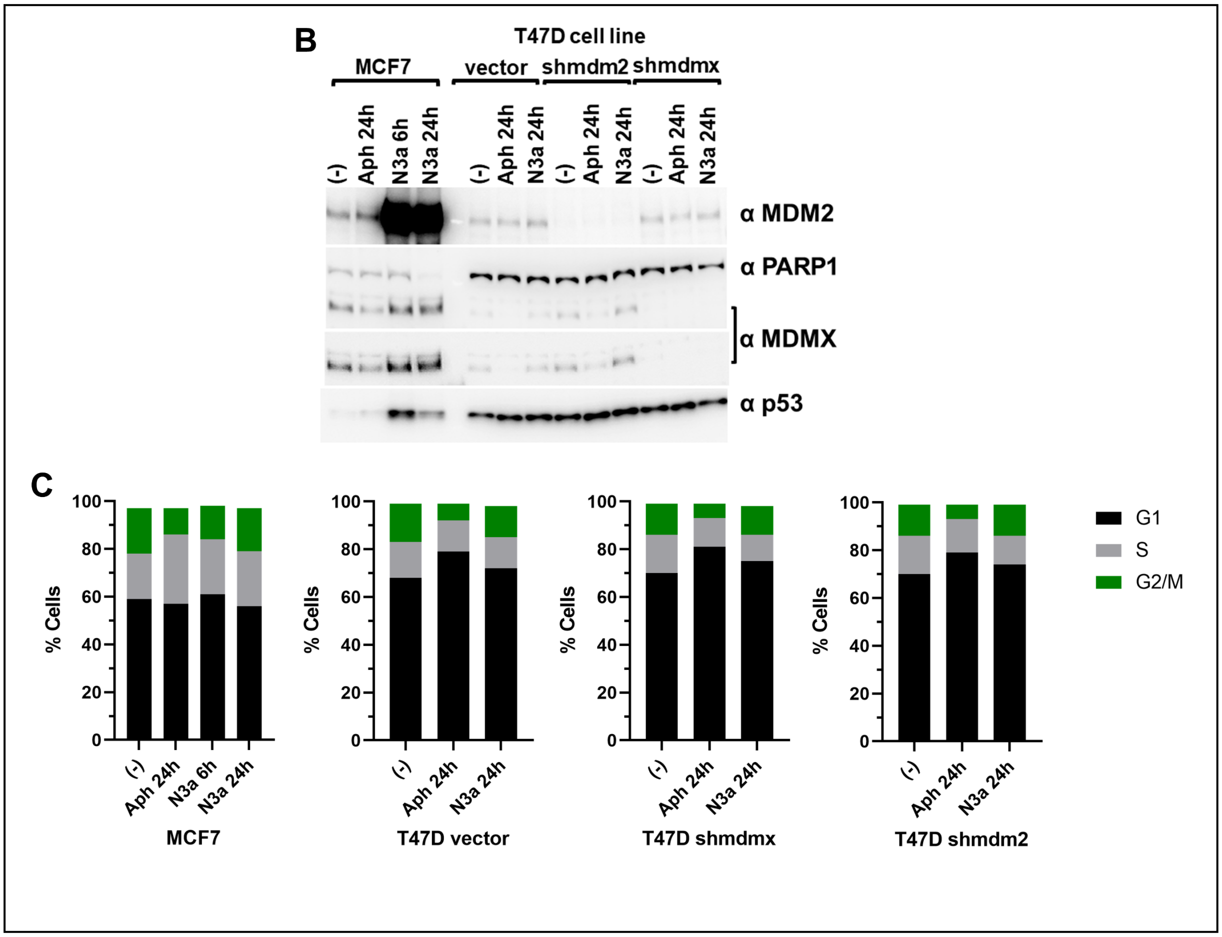


**Panels B and C:** *p53 status of Nutlin 3a-treated breast cancer cells dictates PARP1 stability.*

MCF7 and T47D cell lines were either mock-treated or treated with either 5μM Aphidicolin (Aph) or 10μM Nutlin 3a (N3a) for the indicated time period. The resulting cell populations were analyzed by SDS-PAGE and western blot analysis for the indicated targets using 10μg of each total cell lysate (Panel B), and flow cytometry (Panel C).

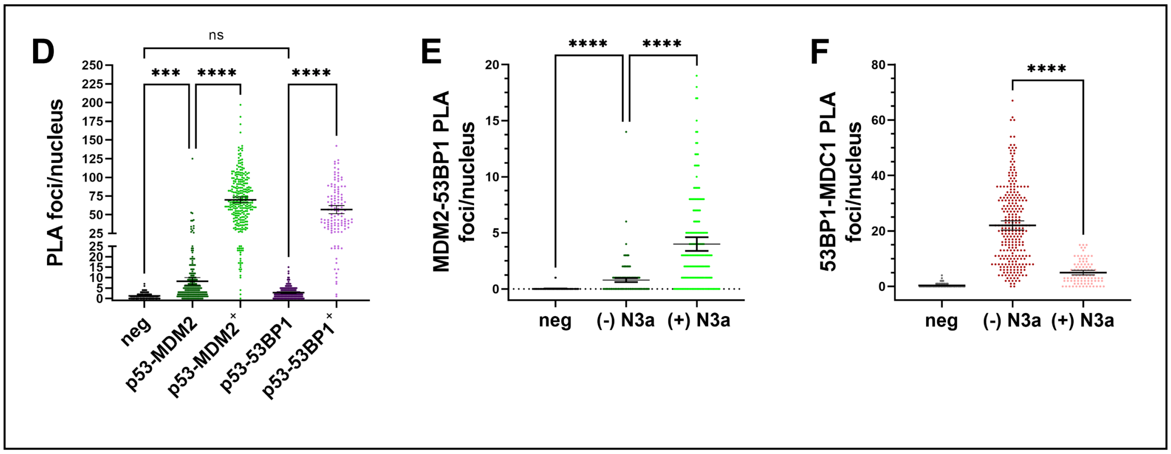


**Panels D, E, and F:** *The influence of Nutlin 3a in MCF7 cells on Proximity Ligation Assay (PLA) interactions between p53, 53BP1, MDM2 and MDC1.*

MCF7 cells treated with 10μM Nutlin 3a for the indicated time periods were analyzed for the p53-53BP1 and the p53-MDM2 PLA interaction (Panel D), the MDM2-53BP1 PLA interaction (Panel E), and the MDC1-53BP1 PLA interaction (Panel F). Confocal images for 3-5 fields for each were acquired, and the number of PLA foci/nucleus from each cell population was determined using Cell Profiler. Graphs and statistical analyses (Kruskal-Wallis) of data from 150-170 cells in panel D, 100-150 cells in panels E, and 150-200 cells F were prepared using Prism. Data representations are mean foci/nucleus with 95% confidence interval from one representative experiment (n=2 for each experiment PLA probe pair); $p < .05$ (*), $p < .01$ (**), $p < .001$ (***), $p < .0001$ (****), $p > .05$ (ns).
